# Run for your life – an integrated virtual tissue platform for incorporating exercise oncology into immunotherapy

**DOI:** 10.1101/2020.09.04.283416

**Authors:** Josua Aponte Serrano, Amit Hagar

## Abstract

The purpose of this paper is to introduce a novel *in silico* platform for simulating early stage solid tumor growth and anti-tumor immune response. We present the model, test the sensitivity and robustness of its parameters, and calibrate it with clinical data from exercise oncology experiments which offer a natural biological backdrop for modulation of anti-tumor immune response. We then perform two virtual experiments with the model that demonstrate its usefulness in guiding pre-clinical and clinical studies of immunotherapy. The first virtual experiment describes the intricate dynamics in the tumor microenvironment between the tumor and the infiltrating immune cells. Such dynamics is difficult to probe during a pre-clinical study as it requires significant redundancy in lab animals and is prohibitively time-consuming and labor-intensive. The result is a series of spatiotemporal snapshots of the tumor and its microenvironment that can serve as a platform to test mechanistic hypotheses on the role and dynamics of different immune cells in anti-tumor immune response. The second virtual experiment shows how dosage and/or frequency of immunotherapy drugs can be optimized based on the aerobic fitness of the patient, so that possible adverse side effects of the treatment can be minimized.

## Introduction

Computational modeling is playing increasingly important roles in advancing a system-level mechanistic understanding of complex interrelated biological processes. In silico simulations guide and underpin experimental and clinical efforts and can advance the knowledge discovery and accelerate therapeutic development at an unprecedented rate. Here we present a computational platform that can interrogate potential mechanisms underlying the effect of aerobic fitness on anti-tumor immune response. These effects, documented in pre-clinical [1,2] and clinical studies [3,4], furnish us with a natural backdrop for probing patient variability and support the inclusion of aerobic fitness as a biological variable in clinical contexts. Doing so may contribute to the personalization of immunotherapy by optimizing dosage and frequency of treatment and by reducing the risk of cardio-toxicity [5] and other adverse side effects.

The model is based on the open source platform of CompuCell (CC3D) which was used in [6] to represent tumor growth as a function of available resources (e.g., glucose). That original model focused solely on differential cell-adhesion and somatic evolution. Since the platform is highly modular, we used the same framework to build a model that includes immune cell types, cytokines, chemokines, and metabolic signals, and employed it to interrogate immune response to tumor progression as a function of aerobic fitness. Our model is a 2D spatiotemporal representation of a cross section in the tumor microenvironment (TME) that allows visualization of the intricate dynamics between the tumor cells and the host immune response. Using data from pre-clinical and clinical studies, we have calibrated the model and estimated the range of the necessary parameters, performed sensitivity analysis, tested the robustness of the fitness parameter, and checked the biological significance of the model by comparing its outcomes to data from clinical and pre-clinical literature, including our own studies. Once it was calibrated, the model was used to test several hypotheses in exercise oncology, and to perform a virtual experiment that probed the effect of aerobic fitness on cardio-toxicity as a potential adverse effect of immunotherapeutic drugs.

Our basics assumption is that aerobic fitness acts as a tumor suppressor through a systemic enhancement of anti-tumor immune response. This systemic effect is a result of metabolic and endocrinal modifications, which can be modulated with chronic exercise training. While the exact mechanisms behind this effect are currently under investigation, documented pre-clinical experiments point at two potential candidates: mechanism (1) increased trafficking of NK cells into the TME, triggered by up-regulation of epinephrine [7,8] and mechanism (2) hypoxia-tolerant suppression of the recruitment of immune inhibitory cells (Tregs) into the TME which increases cytotoxic T lymphocytes (CTLs) efficiency [9,10,11]. In the model presented here we chose to focus on candidate (2),^1^ but the platform can be easily adjusted to incorporate candidate (1) or any other potential mechanism in the future.

## Methods

### Motivation

The clinical data that motivated the model was obtained from a pilot study [12] where recently diagnosed early stage Invasive Ductal Carcinoma patients were subjected to a short submaximal aerobic exercise and were assigned an aerobic score [13]. Tumor size estimated in two time points (the diagnostic mammogram and an earlier mammogram where the radiologist could identify the tumor with hindsight), along with the time between the two mammograms, yielded an estimation of tumor doubling time for each patient. A statistically significant correlation was then detected between the aerobic score and the tumor doubling time: the more aerobically fit were the patients, the longer were their doubling times. Further pre-clinical studies detailed below allowed us to replicate this phenomenon and to interrogate the potential mechanisms underlying it.

### Model description

The model is a spatiotemporal representation of a TME of a solid tumor in its early stages (T0 to T1) that includes key aspects of the interactions between tumor cells, the TME, and the host immune response. In our model, tumor cells adopt four different phenotypes: “oxphos” (performing oxidative phosphorylation), “glycolytic” (elevated glycolysis when the surrounding tissue becomes hypoxic), “necrotic” and “apoptotic”. Tumor cells grow, divide and invade their environment. The growth rate of tumor cells is limited by the availability of oxygen (a field in our model), which cells consume from the environment. In addition to representing the level of tumor immunogenicity (via its effect, mediated by aerobic fitness, on the host immune response), in our model oxygen serves a dual purpose of controlling tumor cell transition from one metabolic phenotype to another (due to oxygenation levels) and tumor growth (due to metabolic resources). As oxygen gets depleted, tumor cells change their metabolic phenotype from “oxphos” to “glycolytic”. Glycolytic tumor cells can change phenotype back to “oxphos” if the oxygenation of the tissue is restored (i.e. the oxygen field strength increases). Otherwise, when oxygen is severely depleted, glycolytic cells become necrotic and die (this phenomenon is typically observed at the tumor core). Glycolytic cells and necrotic cells secrete lactate (chemoattractant *L* in our model) to the TME that serves as a recruiting signal for the tumor promoter cells.

Our model includes two types of immune cells: Tumor suppressors (“CTLs” in our model) and tumor promoters (“Tregs” in our model). “CTLs” are constantly recruited to the tumor site and infiltrate the TME and induce apoptosis in the tumor cells they come into contact with. Upon contact with tumor cells, “CTLs” also release a cytokine signal to the TME (“IFNγ” field), thus attracting other “CTLs” to their vicinity. The acidification of the TME by the glycolytic cells results in recruitment of “Tregs” to the tumor site. These recruited “Tregs” move through the tissue to areas of higher concentration of *L*. “Tregs” inhibit the “CTLs” they come in close proximity to. This inhibition prevents “CTLs” from inducing apoptosis in cancer cells they come into contact with.

We implemented the model in CompuCell3D (CC3D), an open-source modeling environment that allows specification and simulation of multicellular models, diffusing fields and biochemical networks [14]. CC3D simulates spatial dynamics using the Cellular Potts Model, a modeling framework where cells are represented on a lattice and their spatial properties are governed by an effective energy function (See *Supplementary Material*). Spatial dynamics are decided using a Monte Carlo approach, making each independent run stochastic, in which time is measured in Monte Carlo steps (MCS). Our model is simulated over 10! lattice sites representing up to 5×10! individual cells. Diffusion solvers integrate partial differential equations describing the diffusion of oxygen, *L*, and cytokines across the whole simulation domain. The different outcomes of the simulation are dependent on the parameter values associated with aerobic fitness and with the emergent patterns of TME invasion associated with availability of resources and immune response.

### Parameters estimation

Simulation parameters corresponding to the spatial properties of human solid tumor cells, transport of chemicals and rates of immune response were estimated from the literature (See *Supplementary Material*). Each lattice site corresponds to 16*μm*^2^ such that the simulation domain represents a 16*μm*^2^ tissue cross section. We assumed that cancer cells occupy an area of 256*μm*^2^, which is between twice and 3 times the average size of epithelial cells [15]. We assumed that when sufficient resources are available, tumor cells grow and divide every 24 hrs. Conversely, when resources are depleted cells die within 12 hrs, and when “CTLs” induce apoptosis, cells die within 8 hrs. We estimated the infiltration rates of “CTLs” (1 cell every 1.5 hours) and “Tregs” (1 cell every 1 hour) using intramural density data, showing that the “CTL”/“Treg” ratio is 5:1 [16].

The intrinsic random motility and the contact energy were fixed so that tumor cells can detach from each other and invade the surrounding tissue [6]. We assumed that the homeostatic concentration of oxygen in tissue is 4.3×10^-4^ Mol/L [17]. Transport parameters such as oxygen, *L* and IFNγ diffusion coefficients, oxygen uptake rate and *L* secretion rate were estimated from the literature (See *Supplementary Material*). Aerobic fitness was defined as the oxygen concentration threshold at which tumor cells changed from “oxphos” to “glycolytic”. Different populations were defined with respect to different thresholds. The more aerobically fit a virtual subject is, the more tolerant its tissue will be to hypoxia, and as a result, the threshold for the shift from “oxphos” to “glycolytic” is lower.

### Sensitivity and robustness analysis

We used Indiana University’s large-memory computer cluster Carbonate to execute multiple replicates of the simulation [18]. Each replicate is intended to virtually represent individual cancer patient. To generate statistically meaningful results, 200 replicates representing individual patients across 10 aerobic fitness groups were simulated (20 replicates per group). Computational cost, statistical considerations and our own clinical data guided our choice of virtual cohort size: *n=*20 was sufficient to minimize the standard error within groups while allowing a reasonable simulation time, and the aerobic fitness spectrum was chosen to resemble the metric which was introduced in [13] and applied in [12]). Sensitivity analysis was performed by comparing simulation outcomes for different sets of parameters with respect to the final tumor area per aerobic fitness group. The outcomes of the simulation are highly sensitive to key parameters such as the ratio of intratumoral “CTL”/“Treg”, the maximum growth rate of tumor cells and the glycolytic threshold.

Conversely, the outcomes are robust with respect to perturbation of parameters such as cell size or grid size.

### Mechanisms

#### Glycolysis and hypoxia tolerance

The underlying hypothesis of exercise oncology is that aerobic exercise leads to systemic modifications of non-skeletal-muscle tissue. Consistent with several studies that point at hypoxia and elevated glycolysis as a hallmark of solid tumor progression [19,20], here we hypothesize further that aerobic exercise can modify the tissue’s ability to tolerate hypoxia and to degrade HIF1α, a known upstream factor in recruitment of “Tregs” to the TME. To represent this biological feature in our model we introduced the *L* field, secreted by the tumor cells into the TME (Fig. 2). According to our hypothesis, a solid tumor in aerobically fit (sedentary) hosts will generate weak (strong) *L* field depending on that host’s tolerance to hypoxia. This hypothetical differential in turn drives the variation in immune response to the solid tumor. The model presented here thus embodies our working hypothesis (supported by clinical and pre clinical studies; see [21] for an accessible review) that early stage solid tumors of aerobically fit individuals – who have higher tolerance to hypoxia – will go through the shift from the “oxphos” phenotype to the “glycolytic” phenotype in *lower* levels of oxygen in the TME than sedentary individuals, and, as a result, will exhibit *less* glycolysis-initiated immunosuppressive response than similar tumors of sedentary subjects for the same levels of oxygen in the TME.

**Figure 1.**
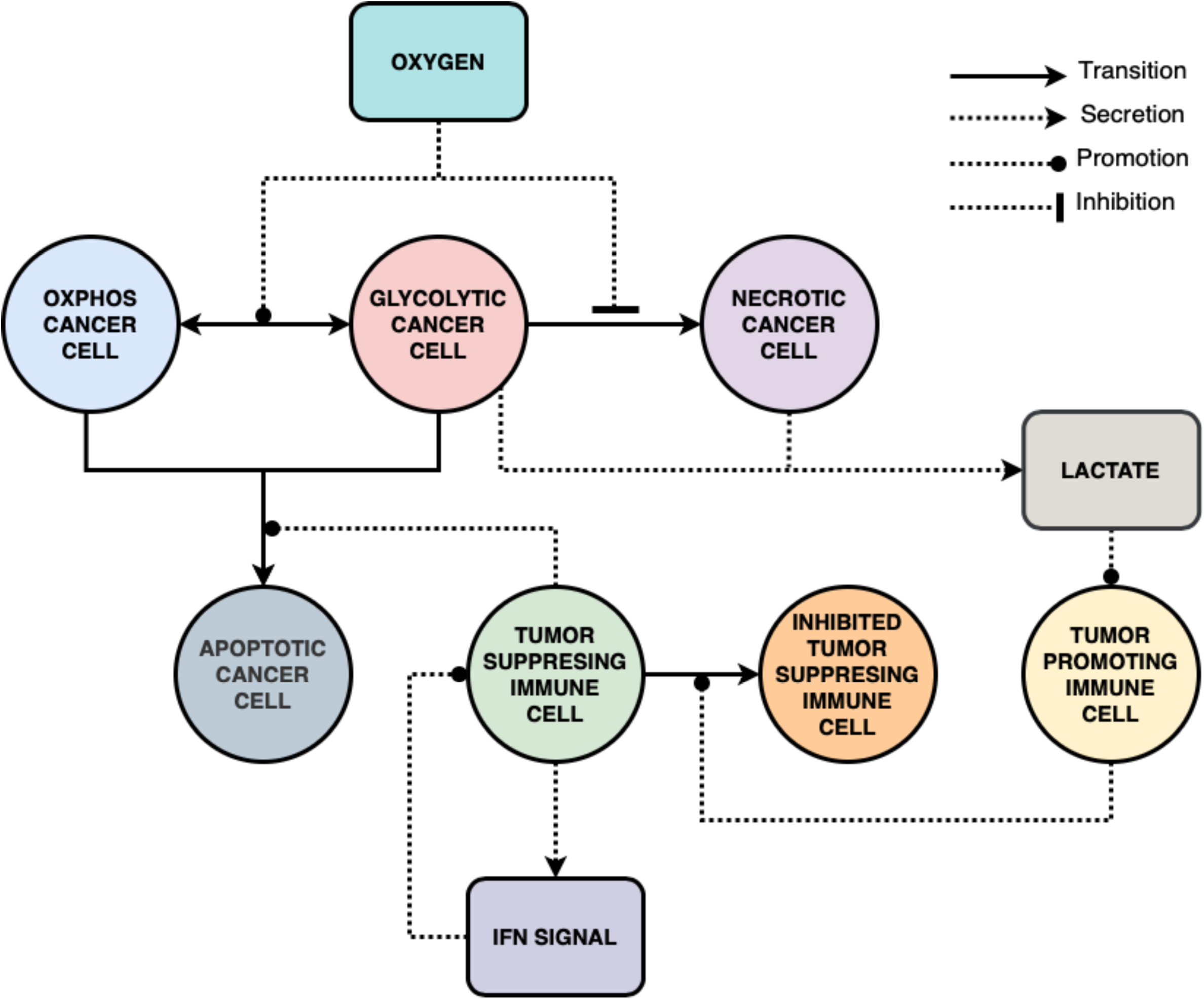
Model Conceptualization. The model simulates the early stage of 2D solid tumor progression from which a growth rate (in terms of tumor area) can be calculated. Once initialized, tumor cells grow in the TME, and with time become more glycolytic, in a rate that depends on the host’s aerobic fitness and tolerance to hypoxia. Tumor cells die through necrosis or apoptosis (lack of oxygen or death by immune response, respectively). Tumor suppressors (“CTLs”) and tumor promoters (“Tregs”) react to cytokine and chemoattractant fields secreted by tumor cells. Tumor cells grow until they saturate the grid.

**Figure 2.**
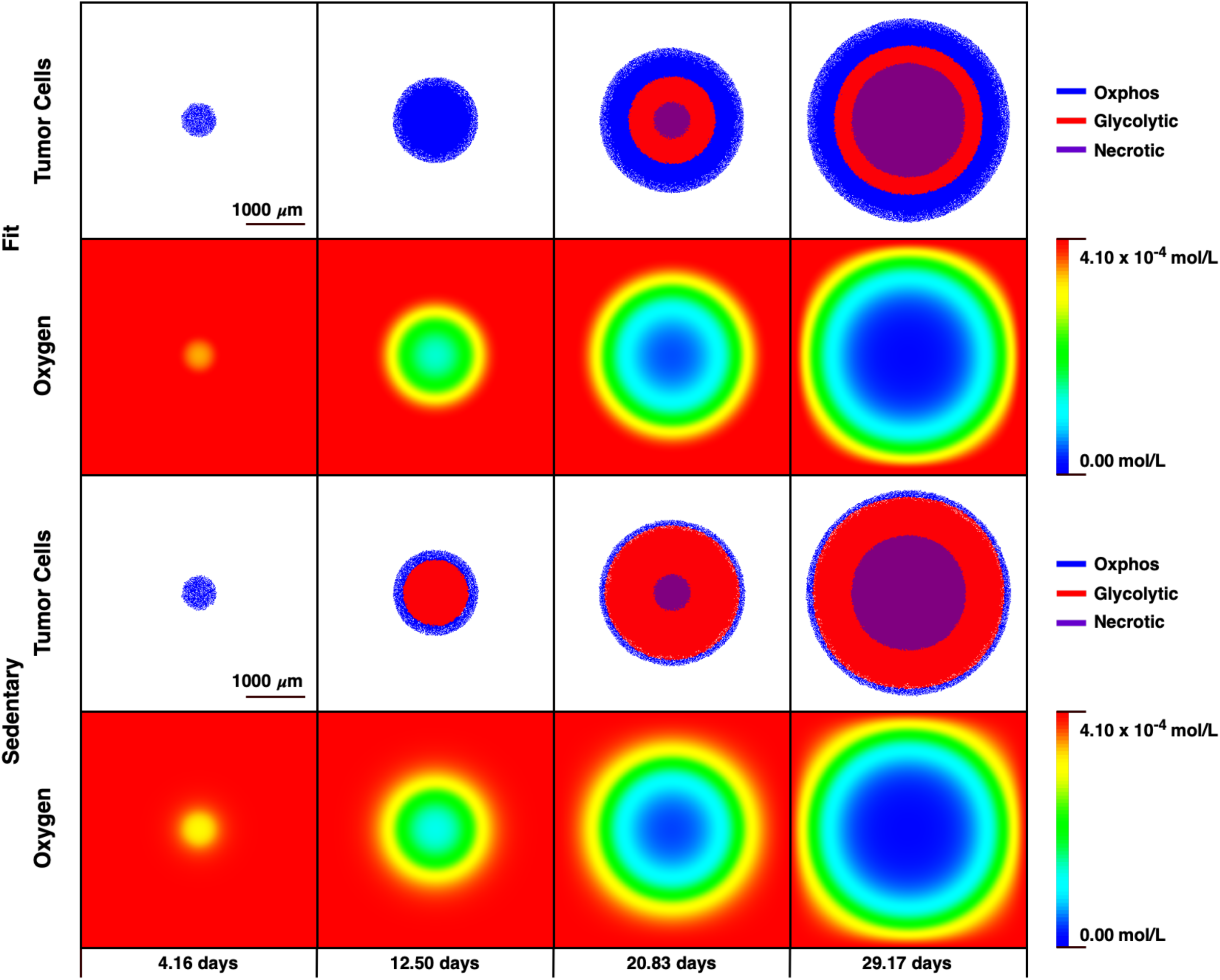
Aerobic fitness modulates hypoxia-tolerance in the TME. While oxygen levels are identical in the two examples above, the two representative TME react to them differently: the more aerobically fit is the host (FIT), the more tolerant to hypoxia its TME is, and as a result, tumor cells are less glycolytic relative to sedentary hosts (SED).

#### Immune suppressors and immune promoters dynamics

Clinical studies have shown that intratumoral CTLs/Treg ratio is a significant prognostic marker for cancer patients [22] and several pre-clinical studies have tied this marker to hypoxic conditions in the TME [10,11]. To represent these biological features in our model we introduced two types of immune cells (immune suppressors, or “CTLs” and immune promoters, or “Tregs”) and implemented two scales of trafficking (Fig. 3). The first is the appearance of these cells in the TME, implemented with different “seeding” rates and densities; the second is movement within the TME, implemented with a chemotaxis mechanism. The seeding rates and densities were calibrated using data on respective densities from hot *vs*. cold tumors in humans [15]. The chemotaxis mechanisms are sensitive to two fields. “CTLs” react to a cytokine secreted by tumor cells killed by other “CTLs” (the “IFNγ” field); “Tregs” react to the *L* field secreted into the TME (the more glycolytic is the tumor, the stronger is the “Treg” recruiting signal).

**Figure 3.**
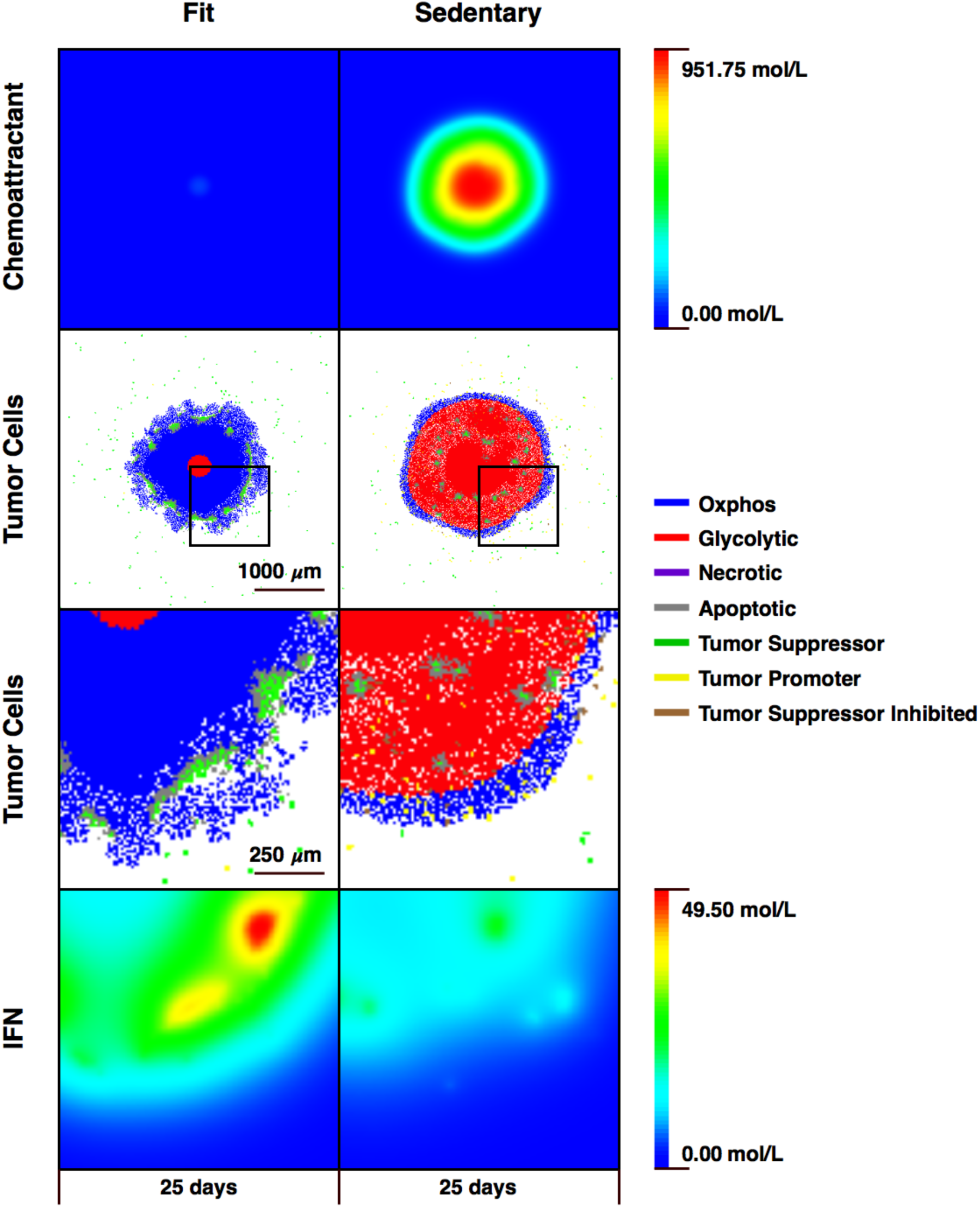
Aerobic fitness modulates anti-tumor immune response. The more aerobically fit is the host, the less glycolytic its tumor cells are relative to a sedentary host. Consequently, recruitment of tumor promoters that can block tumor suppressors is down regulated relative to a sedentary host, and tumor growth will be relatively suppressed. Tumor promoters move towards the tumor along the chemo-attractant gradient that glycolytic tumor cells secrete. Tumor suppressors move towards the tumor along a cytokine gradient (“IFNγ”) that necrotic tumor cells secrete. Once infiltrated into the TME, tumor promoters can inhibit the ability of nearby tumor promoters to kill tumor cells.

### Calibration

#### Effect of aerobic fitness on tumor progression rate

Our model consists of a virtual cohort of 200 virtual subjects divided into 10 aerobic fitness levels. Sensitivity analysis on the aerobic fitness parameter detected its upper and lower bounds below and above which the effects on tumor growth remain constant (where the difference between groups above or below those bounds becomes statistically insignificant at *p*>0.35). Reducing the number of fitness groups *in between* the upper and lower bound can reduce the *p* value between the fitness groups – see *supplemental material* – but won’t change the upper and lower sensitivity bounds (which remain statistically significant at p<0.00001). The model connects variations in fitness levels to variations in anti-tumor immune response and consequently to variations in tumor growth rates. To calibrate it we matched it to clinical results from breast cancer patients (where the aforementioned aerobic score metric was used, and the study relied on hindsight from pre-diagnostic screening mammograms to estimate tumor growth rates in 14 recently diagnosed patients) [12]. The model yielded a classification of distinct aerobic fitness levels, each of which yields a distinct tumor growth curve (Fig. 4). A similar effect of suppression of tumor growth when inoculation followed endurance exercise was qualitatively demonstrated in pre-clinical studies [23,24].

**Figure 4.**
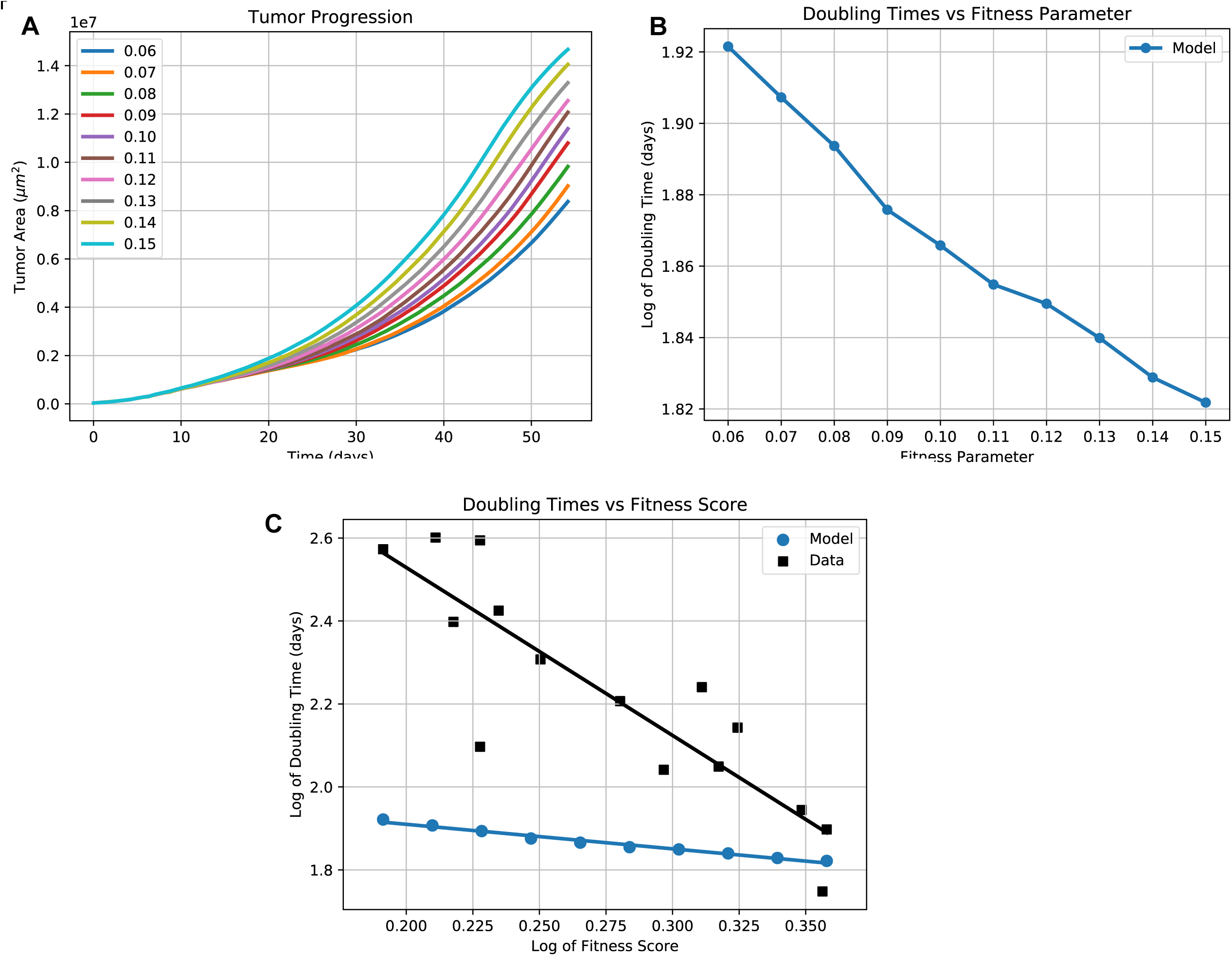
Effect of aerobic fitness on tumor progression rate. The model was run on 200 virtual subjects, divided into 10 distinct aerobic fitness levels, each with 20 subjects. Each fitness level generated an average growth rate (4A). These average growth rates were plotted against the fitness levels on a logarithmic scale (4B). The model behaves qualitatively in accordance with a similar plot of tumor doubling times *vs*. fitness levels from a pilot study in recently diagnosed T1 invasive ductal carcinoma patients (4C) [12]. The comparison between the two correlations (the observed and the mechanistically generated) can be used to further calibrate model parameters (4D).

#### Distribution of tumor doubling time in the population

Studies based on mammography readings [25] have discovered a distribution of tumor growth rates in the population (measured in tumor doubling times). We ran a virtual experiment on our platform with 200 virtual subjects divided uniformly into different aerobic fitness classes and compared the growth rate distribution to the one observed among humans. Both distributions (the virtual and the epidemiological) were lognormal (Fig. 5).

**Figure 5.**
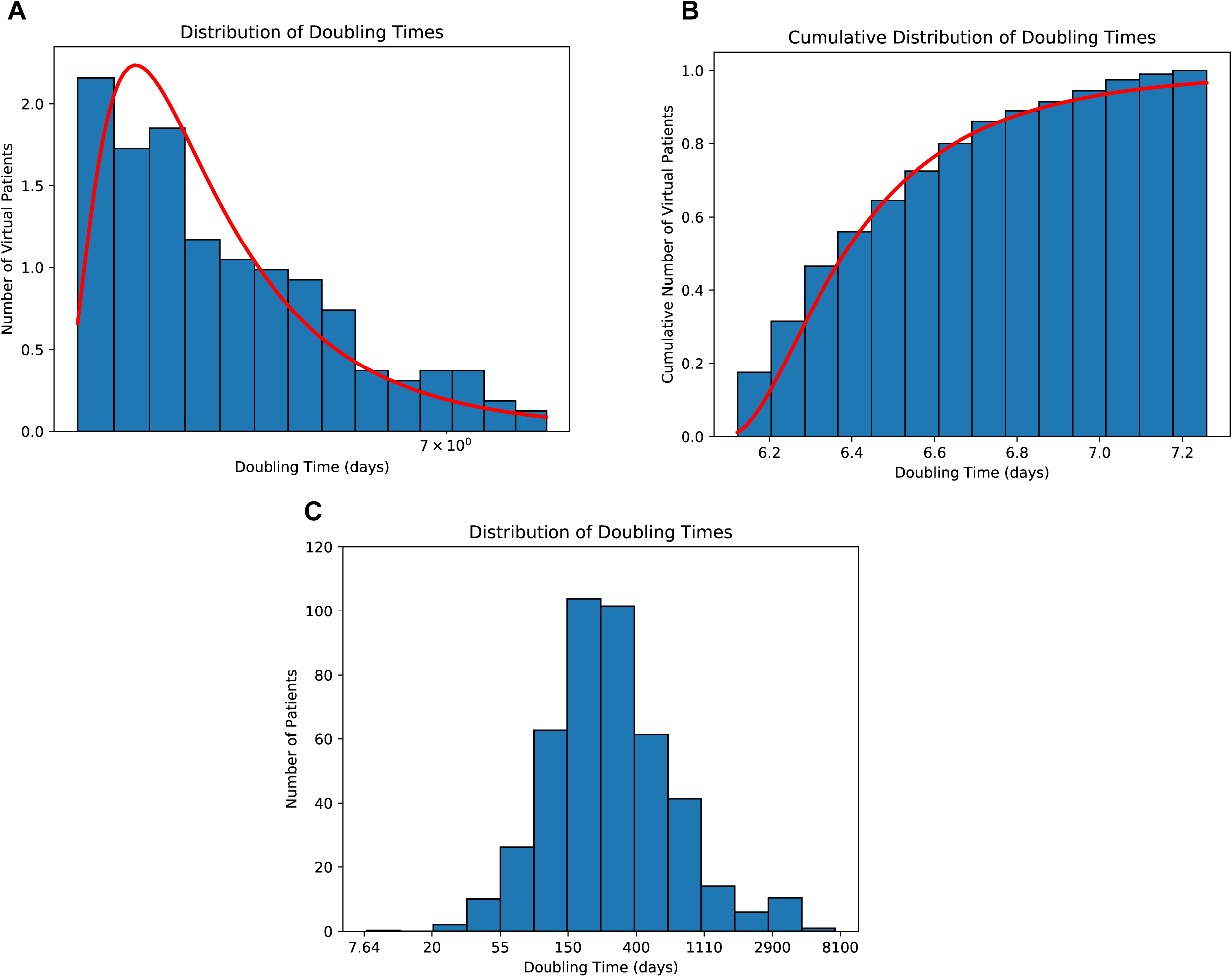
Distribution of tumor doubling time in the population. We ran the model with 200 subjects with random fitness levels. The statistical distribution of growth rates (5A,B) was statistically indistinguishable in a KS test (p>0.29) from a log normal distribution, such as the one observed of growth rates of invasive ductal carcinoma (5C, [25]).

#### Prevalence of clinical tumors in athletes vs. non-athletes

Finally, to identify a fitness threshold that can distinguish in real life aerobically fit from sedentary subjects with the fitness parameter in our model we used epidemiological studies on tumor incidence among athletes and non-athletes [3] (Fig. 6A). The comparison allowed us to designate as “athletes” populations whose fitness parameter value in our model is below 0.060, and sedentary whose fitness parameter value in our model is above 0.150 (the lower this number the higher the aerobic fitness (Fig. 6B)). These values were then matched to the fitness score reported in [21] (where athletes were scored below 0.5 and sedentary above 2 based on their cardiorespiratory fitness (Fig. 4)). Lacking additional granularity of staging variations upon detection in both parameter values groups, we limited this calibration to the identification of a biologically significant spatiotemporal scale for the model: the incidence ratio observed in athletes and non athletes humans is achieved in our model for tumor size of 2.9*mm* and after 13,000 MCS (equivalent to 54 days in real life, and 7 hrs in our simulation) (Fig. 6B). Assuming a clinical (detectable) tumor threshold is 2*mm*, we can use the incidence ratio to compare the scale of our virtual platform to clinical data: our simulated tumors are at least at a scale of ~ 3:2 (thus within the same spatial order of magnitude of a real tumor). Additional data on the staging distribution upon detection in both groups could improve this calibration.

**Figure 6.**
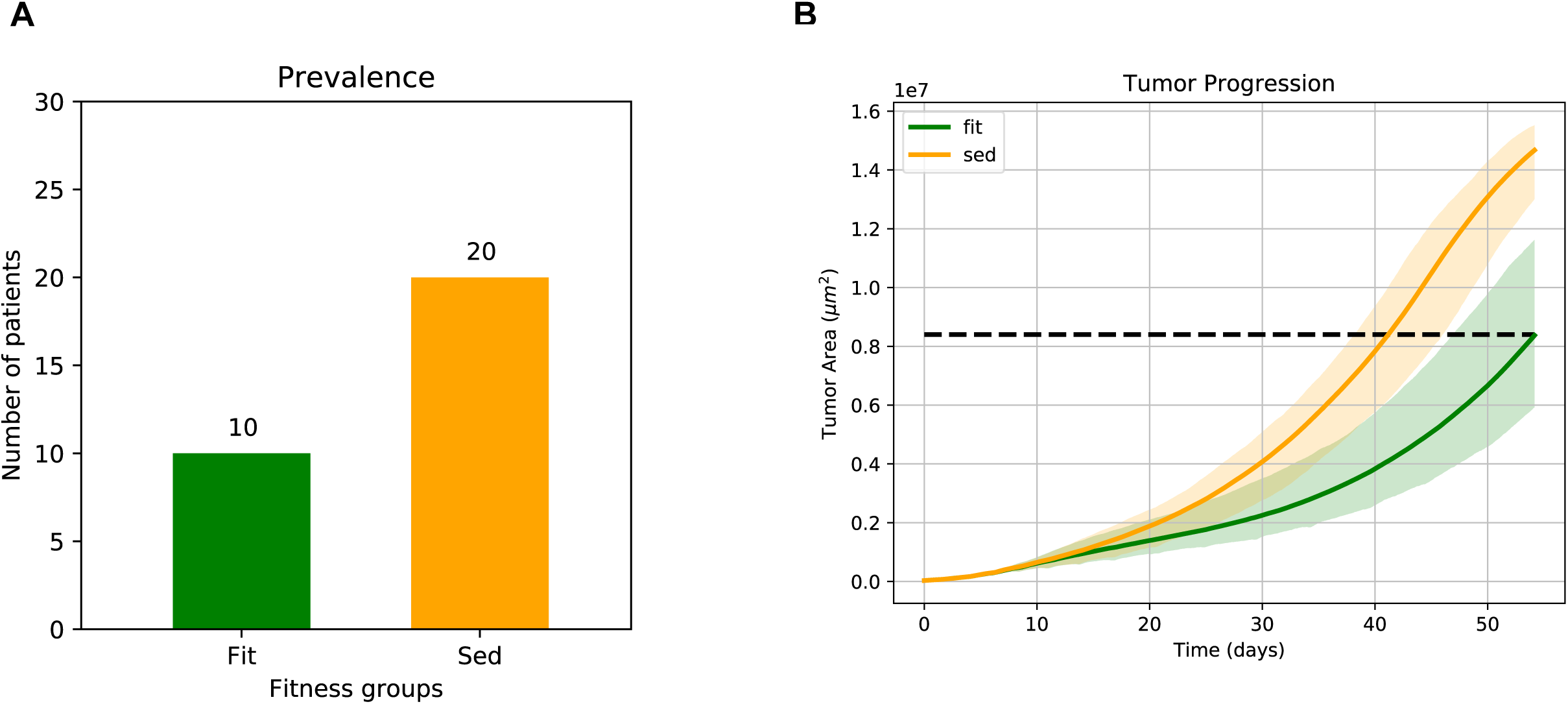
Prevalence of clinical tumors in athletes vs. non-athletes. Epidemiological data shows prevalence of solid tumors in non-athletes to be around twice the prevalence in athletes (6A) [3]. We used this data point to extract a spatiotemporal calibration of the model by running 40 subjects, aerobically fit and sedentary, and identifying the tumor size (in terms of tumor area) and the time after initiation of 200 cells (in model time steps MCS) in which such a prevalence ratio is achieved (6B). The prevalence ratio allows us to impose a spatiotemporal scale on our model (in this case, a scale of 3:2 between model to reality).

## Results

### Time series of anti-tumor immune response in the TME

Our first virtual experiment probes the intricate dynamics in the TME between the tumor and the infiltrating immune cells. Such dynamics is impossible to probe in humans, and is hard to observe during a pre-clinical study as it requires significant of redundancy in lab animals (so to achieve high resolution with statistical significance for successive end-points during the experiment), prohibiting such time and labor intensive studies. Our simulation generates, with no physical cost, a time series of spatiotemporal snapshots of the TME (Fig. 7) that can serve as a platform to test several mechanistic hypotheses on the role and dynamics of different immune cells in ant-tumor immune response, by comparing it to immunohistochemistry slides from different stages of tumor development. While here we focused only on two types of immune cells (“CTLs” and “Tregs”), and two types of signaling fields (“IFNγ” and “chemoattractant”, or *L*), the platform is modular and can incorporate many more cells and fields (hence more pre-clinical end points) with relatively small modifications.

**Figure 7.**
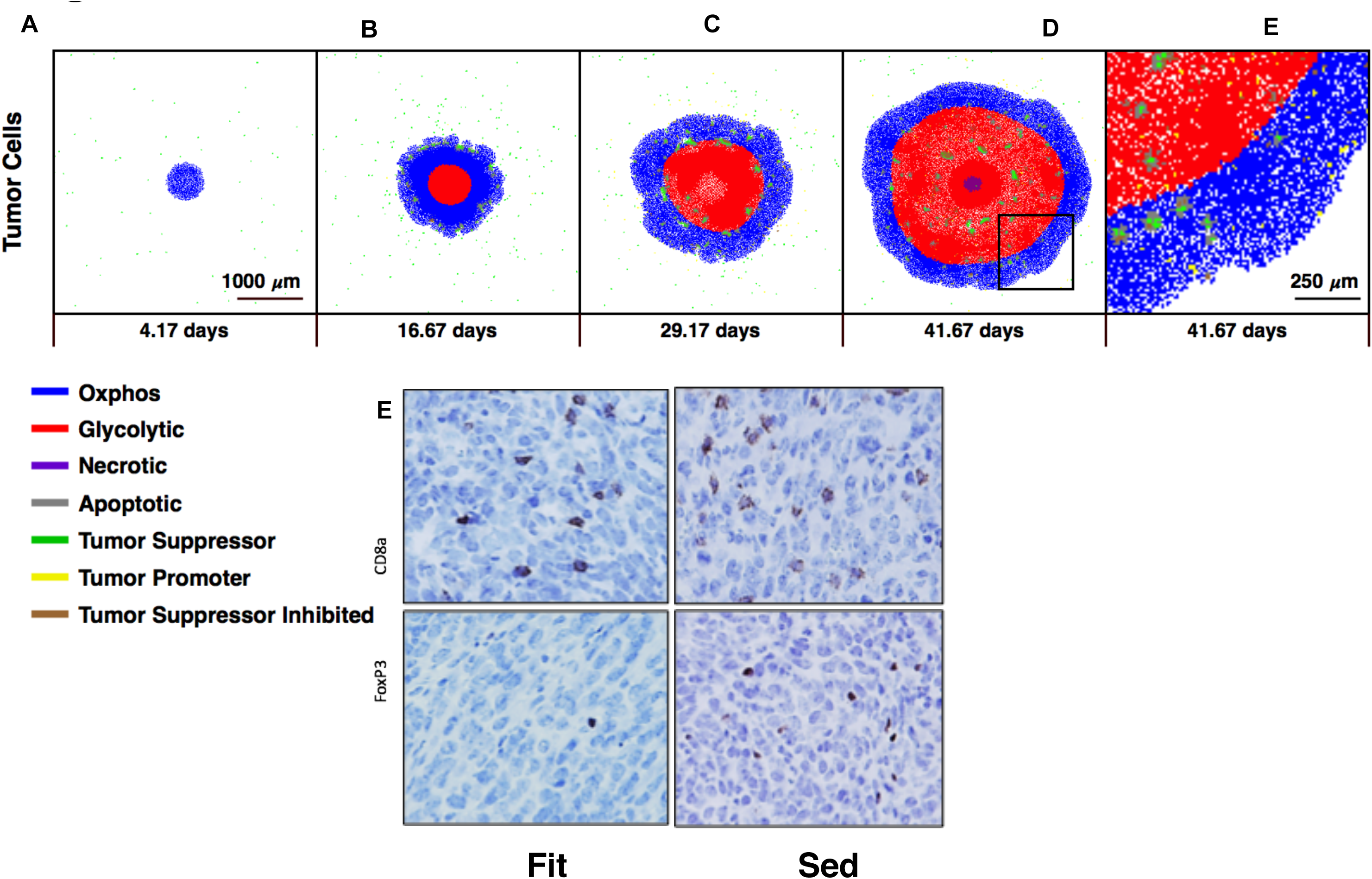
Time series of TME sections in early stage progression of a solid tumor. To probe the intricate dynamics of anti-tumor immune response in the early stages of a solid tumor progression, the model can yield an observation window into the TME in different stages of growth (7A-D), and can be used to test competing hypotheses on tumor immune cells population dynamics by comparing these snapshots to real life immunohistochemistry end points (7E, [23]), where cross sections from exercised (“FIT”) and sedentary (“SED”) mice show different intratumoral CD8^+^/ CD4^+^FOXP3^+^ ratios. Additional plots in *Supplemental Material* (Fig. S3) show the potential of the model to generate quantitative analysis for TME markers which can be compared to desired pre-clinical end points.

### Incorporating aerobic fitness into the personalization of immunotherapy

While showing remarkable success in some patients, immunotherapy treatments can lead to severe autoimmune adverse effects such as myocarditis, pericardial diseases, and vasculitis, including temporal-arteritis and vision loss [5]. To mitigate those, careful dosing is essential. If our hypothesis on aerobic fitness as a biological variable is correct (and there is some support for it from pre-clinical studies on the combination of Immune Checkpoint Inhibitors (ICI) with aerobic exercise [24,26] and from small pre-treatment exercise intervention in humans [27], and our own clinical pilot study [12], see Fig. 8G), aerobically fit patients may require lower dosage of ICI than sedentary patients, which may lead to personalization of treatment and reduction of adverse effects. To test this hypothesis we implemented ICI in our model as an increased efficacy of “CTLs” killing, by limiting the inhibitory radius of the “Trges”. Cytotoxicity was then quantified with the “IFNγ” field, where the probability of an adverse effect [28] increases with that field’s strength. Performing a virtual experiment on both aerobically fit and sedentary virtual subject populations treated with ICI, the model shows how, without a mitigated dosage, aerobically fit subjects are more prone to adverse effects than their sedentary counterparts (Fig. 8A,B). Conversely, lowering the dosage of ICI for aerobically fit patients can achieve the same reduction of tumor growth relative to their sedentary counterparts but with a lower probability for adverse effects (Fig. 8C,D). In order to translate this result to a clinical setting as in 8E,F, future studies should identify potential markers for aerobic fitness with which such personalization can be accomplished and a clinical study could be performed to test the appropriate dosage of ICI in sedentary *vs* aerobically fit patients.

**Figure 8.**
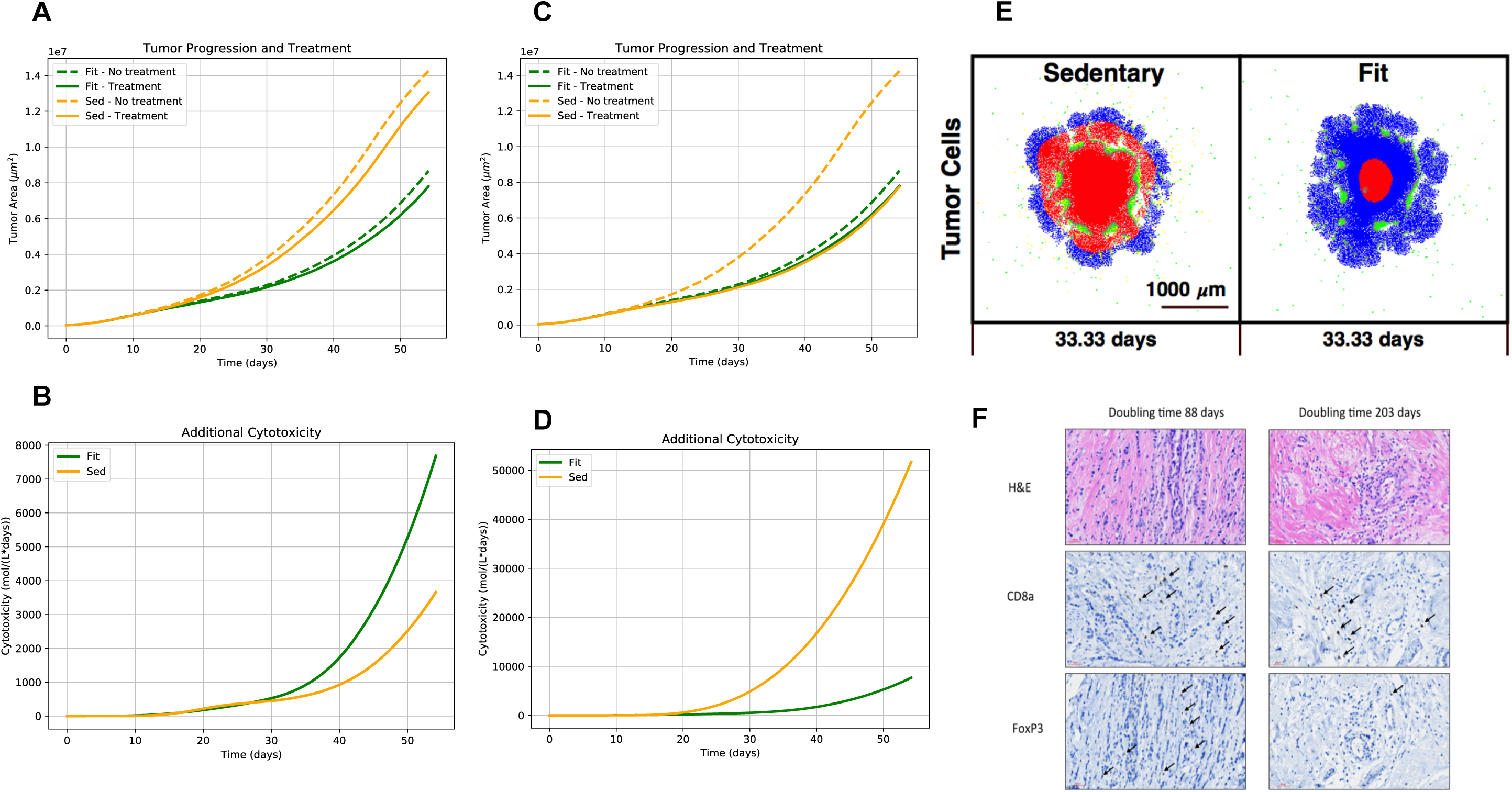
Precision immunotherapy. Aerobically fit patients may require smaller dosage of ICI than sedentary patients, which may lead to personalization of treatment and reduction of adverse effects. Without a mitigated dosage, aerobically fit subjects are more prone to ICI adverse effects than their sedentary counterparts (8A,B). Lowering the dosage of ICI for aerobically fit patients relative to their sedentary counterparts can achieve the same reduction in tumor growth (8C) but with a lower added toxicity hence lower probability for adverse effects (8D). As a result of the ICI, the two tumors in 8E (sedentary and fit hosts), treated with high and low dosage, respectively, are of the same size, regardless of their initial immunogenicity. IHC of fast and slow growing Invasive Ductal Carcinomas in human females from the study reported in [12] show respectively lower and higher ratios of CD4^+^FOXP3^+^ to CD8^+^ T cells (8F).

## Discussion

### What we have shown

After calibrating our model with clinical and epidemiological data, we performed two virtual experiments that showcase the potential usefulness of the platform as a tool for improving pre-clinical and clinical studies. We have shown how to generate a time series of TME snapshots during anti-tumor immune response, and how to personalize dosing of ICI for aerobically fit patients in order to lower the risk of adverse effects. Further collaboration with cancer biologists on *in vivo* studies and clinicians on human studies would allow us to implement the platform for the purpose of improving *in vivo* experimental design and personalization of clinical outcomes. We have deposited the model on a 3^rd^ party open source repository (GitHub), so that users can download it and modify it for research purposes.

### Justification

Endurance exercise has been shown to be a systemic modulator of metabolic and endocrinal activities, and, through these, a modulator of immune competence and a natural element in cancer prevention. Here we propose to treat aerobic fitness as a biological variable that can be incorporated into cancer (immuno)therapy and improve personalization of treatment.

The exact underlying mechanisms behind the suppressive effects of aerobic exercise on early tumor progression are currently unknown. Several pre-clinical studies have narrowed the possibility space down to two main hypotheses [29]. The first involves exercise-induced up-regulation of epinephrine that mobilizes NK cells into the TME [7,8] together with increased trans-signaling of IL-6 [30,31] as a re-distributing factor (increased adhesion, infiltration and activation). However, since these effects were induced on mice by exposing them to voluntary running, their human relevance is suspect [32]: voluntary running in mice mimics high intensity interval training (HIIT) in humans, and no human, even an elite athlete, can endure the “dosage” level of HIIT exhibited in those studies [33,34]. Since there are currently no experiments testing this hypothesis with more human-relevant, lower HIIT “dosage” (in which the voluntary wheel is activated only for a short period per day), and since our ultimate goal is to use the model for personalization of patient outcomes, our model focuses on the second candidate the human-relevance of which is more significant.

This second hypothesis, the one that underlies the model presented here, connects exercise-induced increased hypoxia-tolerance to more efficient anti-tumor immune response, and requires chronic endurance training (CET) which can be achieved in pre-clinical exercise oncology with forced running wheels [23,32]. The idea here is that CET induces hypoxia tolerance in the skeletal muscles and in other tissues, and as a result, TMEs are more susceptible to the degradation of HIF1α [35]. This degradation is an upstream factor in a signaling cascade leading to increased anti-tumor immune efficiency, as HIF1α is known to recruit, via myokine signaling [10,11,36], Trges into the tumor micro-environment, which suppress CTLs [9]. Our pre-clinical study detected a twofold decrease in intratumoral Tregs/CTLs ratio in exercised mice relative to their sedentary counterparts [23]. Support for this second potential mechanism in humans that correlates CET with reduced intratumoral Treg density comes also from a small pilot study in which recently diagnosed breast cancer patients were subject to a short exercise session *before* treatment, and their excised tumor presented with threefold fewer intratumoral Tregs [27]

Attempts to utilize aerobic fitness as a predictor for patient outcomes are not new. For example, frailty indices (which include aerobic fitness, or lack thereof, as one of their components) have been recently used to predict adverse health outcomes in cancer patients post surgery or chemotherapy [37,38].^2^ We have not investigated the possible connection between this phenomenon and the effects of aerobic exercise on tumor growth, but we believe that as our understanding of the mechanisms that underlie these effects will increase, more connections between aging and cancer will be unearthed and will be explained with common pathways.

### Limitations

The model presented here includes only two types of immune cells: tumor suppressors and immune inhibitors (or tumor promoters). We deliberately chose to start from the simplest model under the assumption that any introduction of additional immune cells could increase the ability of the model to replicate observed phenomena, rather than decrease it. Given our initial success, work is currently underway to include additional types of immune cells in future versions of the model.

Our platform can perform virtual experiments with no wet-lab or clinical costs, and is proposed here as a tool for pre-clinical and clinical researchers. The tool is limited in several ways. First, to obtain simulation results in a reasonable time we must limit the computational cost. Consequently, our grid size is currently bounded by 5×10^4^ cells. This size allows the simulation to be sensitive to spatiotemporal and stochastic features of the dynamics; increasing it will incur computational cost, but given the running time of the current simulation (7 hrs per one virtual subject), it shouldn’t be prohibitive on a multi-core platform where subjects are run in parallel. Second, the same concerns about computational cost limit us to two-dimensional simulations. Specific circumstances may require scaling up to 3D (e.g., angiogenesis), which can be done on segments of the grid to reduce complexity. For most clinical end-points, however, a cross section of the TME may be a good approximation. Third, here we introduced only two types of immune cells and three types of fields. Increasing the number of members of both sets will, again, add complexity (and computational cost) to the model. From our experience, however, a direct dialogue between model developers and clinicians may help optimize the platform for each specific usage so that the computational cost won’t become prohibitive. Finally, note that we deliberately calibrated the model solely with quantitative clinical and epidemiological data, and limited the usage of pre-clinical studies to qualitatively probing mechanistic hypotheses. We did so because our platform faces a challenge prevalent in biomedical research in general, namely, spatiotemporal scaling between mice models and human models. Acknowledging this challenge, we believe we are better equipped to meet it: once the model is separately calibrated for each modality, it allows us to virtually compare time series from pre-clinical simulations to observed clinical end-points (a process that is hard to mimic in real-life experiments) and help to characterize the scaling between murine and human immunological clocks.

### Insights and future directions

Solid tumor progression is an intricate process that can be characterized with population dynamics where tumor cells and immune cells are intertwined in complex signaling networks. As such it is sensitive to spatiotemporal factors which are difficult to incorporate into current machine-learning-based bioinformatics platforms. Causal multi-scale modeling is a viable alternative that can mitigate this shortcoming; our *in-silico* modeling platform of anti-tumor immune response and early stage solid tumor growth presented here is an example. The long-term goal of all such modeling, which can only show that a specific underlying mechanism is *sufficient* to explain the observed phenomena, is the development of quantitative, predictive models based on clinical and experimental data which will have a positive impact on patient outcomes through improved patient-specific treatment regimes.

Perhaps more than any other therapy, cancer immunotherapy is particularly sensitive to timing [39]. Efforts invested in examination of combination dose scheduling can yield qualitatively significant returns in terms of improved efficacy and decreased toxicity; all without necessitating regulatory approval. Our *in silico* platform is a safe playground for such experimentation in dosage scheduling and frequency, as it can easily allow modulation of duration and timing of activation signaling to achieve the most effective treatment, thus ruling out extreme scenarios and refocusing the researcher on an optimal treatment window, while harmlessly testing different dosage regimes, frequency and duration.

Finally, our platform can easily incorporate and test combination of different types of immunotherapy with other standard-of-care therapies [40] and probe potential synergistic effects. For example, since aerobic exercise promotes oxygenation, it can mimic the effects of Anti Angiogenic Therapy, where different aerobic fitness levels can be calibrated to represent different dosage of such a therapy.

To conclude, we presented a small sample of what our integrated platform can offer. To fully achieve its potential, it requires careful calibration with clinical data and pre-clinical experiments. To this end, we hope to start a dialogue with interested cancer immunotherapy clinicians and pre-clinical cancer researchers who are open to incorporating *in silico* modeling into their experimental portfolio.

## Abbreviations

CC3D: CompuCell 3D
CET: chronic endurance training
CPM: cellular Potts model
CTL: cytotoxic T lymphocytes
HIIT: high intensity interval training
ICI: immune checkpoint inhibitors
MCS: Monte Carlo steps
TME: tumor microenvironment

## Declarations

### Ethics approval

N/A

### Consent to participate

N/A

### Consent for publication

N/A

### Availability of data and material

All data was retrieved from published literature at PUBMED

### Code availability

Code is available from JAS upon request

### Authors’ contributions

Conceptualization: AH. Model conceptualization: AH & JAS. Coding and Running Simulations: JAS; Manuscript writing and editing: AH & JAS.

## Acknowledgments

The authors acknowledge the Indiana University Pervasive Technology Institute for providing HPC (Carbonate) resources that have contributed to the research results reported here. URL: https://pti.iu.edu/. We thank James Glazier (IUB) for useful discussion.

## Footnotes

1 Our choice is motivated by the methodological problems that surround the pre-clinical studies behind mechanism (1) which in turn make its exercise “dosage” less human-relevant than the one induced in mechanism (2). See Discussion.

2 We thank an anonymous referee for this comment.

