## Supplementary Material for "Run for your life – an integrated virtual tissue platform for incorporating exercise oncology into immunotherapy"

### Biological and Mathematical Model

#### Cell Spatial Dynamics

We model cellular spatial dynamics using the Cellular Potts Model (CPM or Glazier-Graner-Hogeweg model) which represents generalized cells as occupying sets of voxels on a fixed lattice. These generalized cells can represent biological cells, cellular subcomponents or extracellular domains. Each voxels in the lattice has a position $x$ and an index associated with the generalized cell occupying that voxel $\sigma(x)$. To represent distinct phenotypic states, generalized cells $\sigma$ are assigned a cell type $\tau(\sigma)$.

In the absence of external stimuli, cytoskeletal membrane fluctuations and differential adhesion to the extracellular matrix drive migration of biological cells in a random-walk pattern. In this model we assumed that tumor cells invade the tumor microenvironment (TME) following a random walk. Random cell motility is stimulated by stochastic exchange of voxels occupied by each generalized cell. Other spatial properties and behaviors are modeled by associating effective energy terms with generalized cell properties such as size and shape constraints (volume, surface) and behaviors such as mechanical interactions (cell adhesion) and directed motion (chemotaxis).

The configuration of the lattice evolves to minimize the system's effective energy:

$$H = \sum_{x}\sum_{y}^{N} J(\sigma(x), \sigma(y)) \left(1 - \delta_{\sigma(x),\, \sigma(y)}\right) + \sum_{\sigma} \lambda_{vol}(v(\sigma) - V(\sigma))^2 + \sum_{\sigma} \lambda_{sur}(s(\sigma) - S(\sigma))^2 \quad (1)$$

The first term models cell adhesion: $N(y)$ is the neighborhood of site $x$, $\delta_{i,j}$ is the Kronecker-delta function, the term $(1 - \delta_{\sigma(x),\, \sigma(y)})$ enforces counting energies between voxels belonging to a different cell and $J(\tau(\sigma(x)),\ \tau(\sigma(y)))$ is the effective contact energy per unit surface area between cells $\sigma(x)$ and $\sigma(y)$. The next two terms model cell volume and surface area as quadratic constraints: $\lambda_{vol}$, $\lambda_{sur}$ denote the strength of, the constraints $v(\sigma)$, $s(\sigma)$ denote the current volume and surface area and $V(\sigma)$, $S(\sigma)$ denote the target volume and target surface area of cell $\sigma$. The values of these constraints can be assigned by individual cells or to groups of cells depending on their "cell type" (discussed below).

The lattice configuration evolves by voxel copy attempt. A target voxel $x_i$ and a neighboring source voxel $x_j$ are randomly selected. If different cells occupy these voxels, the energy change ($\Delta H$) associated with updating the generalized cell at $x_i$ with the one occupying $x_j$ is evaluated. The probability of accepting the voxel copy attempt is given by a Boltzmann acceptance function:

$$\Pr\left(\sigma(x_j) \to \sigma(x_i)\right) = 1 \ \ for\ \Delta H \le 0\ ,\ e^{\frac{\Delta H}{T}} \ \ for\ \Delta H > 0 \ (2)$$

where $\Delta H$ is the change in the systems effective energy from the voxel copy attempt and $T$ is the amplitude of the cell-membrane fluctuations leading to random motility. The intrinsic simulation unit is a Monte Carlo step corresponding to a series of voxel copy attempts.

Model specification and simulations were performed using CompuCell3D (CC3D), an open-source multicellular modeling environment (www.compucell3d.org). CC3D allows rapid specification of multiscale models that integrate spatial cellular dynamics, diffusion of chemical fields and dynamics of biochemical networks using Python. CompuCell3D simulation can be executed on Windows, Mac and Linux platforms and supports cluster execution for parameter exploration.

Parameter Estimation. Following Maciej et al [6], we adopted a voxel length of 4 $\mu m$, such that the each voxel represents an area of 16 $\mu m^2$. The total simulation domain consists of 1000x1000 voxels, corresponding to a tissue cross-section of 16 $mm^2$. The initial tumor cell target volume is 256 $\mu m^2$ and the immune cell target volume is 384 $\mu m^2$. The initial tumor cell surface is 64 um and the immune cell target surface is 78.4 um.The cell surface is calculated from the cell volume assuming that the area occupied by cells is a square. Note that 'cell volume' and 'cell surface' are CC3D-specific cell properties denoting the number of voxels occupied by each cell and the number of outer voxels of each cell respectively. As such, they correspond to the cross-sectional area and the perimeter of cells. The MCS to time conversation is 6 mins. This estimate is based on experimental tumor cell migration speed [6]. For the contact energies, we assigned values that result in a highly negative surface tension between tumor cells and the surrounding general medium such that tumor cells invade the surrounding tissue. These parameters are CC3D-specific and the values are shown in Table 1.

**Cell Types**

'Cell types' are CC3D-specific attributes that allow to classify individual cells based on different phenotypes and behaviors. We represented the different population of cells in the TME by including four types of tumor (*OXPHOS, Glycolytic, Necrotic and Apoptotic*) cells and two types of immune cells (*Tumor Suppressors and Tumor Promoters*). *OXPHOS* represents tumor cells that mostly rely on oxidative phosphorylation as their main metabolic pathway. *Glycolytic* represents tumor cells that mostly rely on glycolysis as their main metabolic pathway. OXPHOS and glycolytic tumor cells grow and divide at the same rate based on the availability of metabolic resources. OXPHOS cells rapidly consume oxygen from the TME while glycolytic cells produce pro and anti-inflammatory signals that recruit immune cells. *Necrotic* represents tumor cells at the necrotic core of the tumor dying due to lack of metabolic resources. Necrotic cells also produce proinflammatory and anti-inflammatory signals. *Apoptotic* represents tumor cells undergoing apoptosis due to immune cell-mediated cytotoxicity. Apoptotic tumor cells shrink and die. *Tumor Suppressors* represent cytotoxic immune cells that can induce apoptosis in the tumor cells they come into contact with. The cytotoxic mechanism is described in *Immune Cell Cytotoxicity*. Tumor suppressors have two states: active or inhibited. 'Active tumor suppressors' are capable of inducing apoptosis whereas 'Inhibited tumor suppressors' are not. *Tumor Promoters* immune cells represent the anti-inflammatory response of the immune system by inducing the transition of tumor suppressors from active to inhibited.

**Chemical Fields and Diffusion**

***Oxygen Transport.*** Although tumor growth in vivo depends on multiple chemical substances (including glucose, growth factors, fatty acids) we introduce the chemical field oxygen ($O$) to represent both tissue oxygenation and availability of other metabolic resources. The change in concentration of $O$ is calculated by solving the reaction-diffusion equation at each location in the simulation domain:

$$\frac{\partial O(x)}{\partial t} = D_O \nabla^2 O(x) - d_{O-OXPHOS} O(x) - d_O\, O(x) + P_{O-Medium}(x) \quad (3)$$

where $D_O$ is the diffusion coefficient of oxygen, $d_{O-OXPHOS}$ is the decay rate of oxygen inside OXPHOS cells (accounting for the oxygen consumption by cells relaying in oxidative phosphorylation), $d_O$ is the global decay rate of oxygen inside all other cells (accounting for consumption and decay of oxygen in general tissue) and $P_{O-Medium}$ is the production rate of oxygen by medium (representing oxygenation from blood vessels).

Parameter estimation. Based on *in vitro* assays, we assumed that the homeostatic oxygen tension of tissue (in the absence of tumor cells) is 4.30 x 10 $^{-4}$ mol/L [15]. We followed the same method as Maciek et al [6], to determine the rate of oxygen production by Medium by assuming that in real tissue stromal cells consume oxygen at a rate 2.00 x 10 $^{-17}$ mol/(cell*s) and that stromal cells occupy 20% of the TME [A]. To maintain a steady state concentration of the production rate of oxygen by Medium is 8.00 x 10 $^{-16}$ mol/s and the global decay rate of Oxygen is 0.01 s $^{-1}$. We assumed that at the steady state levels, tumor cells consume oxygen at a rate 6.00 x 10 $^{-17}$ mol/(cell*s), 3 times faster than stromal cells because of higher metabolic demands. The diffusion coefficient of oxygen in water is 1460.0 $\mu m^2/s$ [B].

***Chemoattractant Transport.*** We introduced the chemical field Chemoattractant ($L$) to represent the signaling cascade that starts with HIF1a stabilization, higher ratio of glycolysis to OXPHOS, lactate secretion and production of chemokines such as CCL-28 that recruit Tregs into the TME. (see [10,11,35] in the main article). The change in concentration of $L$ is calculated by solving the reaction-diffusion equation at each location in the simulation domain:

$$\frac{\partial L(x)}{\partial t} = D_L \nabla^2 L(x) - d_L L(x) + P_{L-Gly}(x) \quad (4)$$

where $D_L$ is the diffusion coefficient of the chemoattractant, $d_L$ is the decay rate of the chemoattractant in the TME and $P_{L-Gly}$ is the production rate of chemoattractant by glycolytic cells. Although *in vivo*, every cell produces lactate as a byproduct of their metabolism, our chemical field $L$ represents the excess of lactate produced by cells with a higher ratio of glycolysis to OXPHOS (which, according to our hypothesis is modulated by aerobic fitness).

Parameter Estimation. The diffusion coefficient of the chemoattractant is in same order of magnitude of the diffusion coefficient of glucose: 0.1 $\mu m^2/s$ [C]. We assumed that the chemoattractant signal is short ranged, such that the diffusion length of the signal is 3 cell diameters (48 $\mu m$). Based on this assumption, we estimated the decay rate of the chemoattractant to be: 1.0 x 10 $^{-7}$ s $^{-1}$. Glycolytic cells produce chemoattractant at the same rate cells produce lactate *in vitro*: 7.52 x 10 $^{-17}$ mol/(cell*s) [D].

***IFN-γ.*** Active immune cells relay information and recruit more immune cells by secreting a variety of chemokines. We simplified the complexity of immune cell signaling by introducing a single chemical field IFN-γ ($IFN$). The change in concentration of $IFN$ is calculated by solving the reaction-diffusion equation at each location in the simulation domain:

$$\frac{\partial IFN(x)}{\partial t} = D_{IFN} \nabla^2 IFN(x) - d_{IFN} IFN(x) + P_{IFN-Immune}(x) \quad (5)$$

where $D_{IFN}$ is the diffusion coefficient of IFN, $d_{IFN}$ is the decay rate of IFN in the TME and $P_{IFN-Immune}$ is the production rate of IFN by tumor suppressor cells.

Parameter Estimation. Given we are modeling the TME, in this short scale we can assume the same transport coefficients for the IFN field and the chemoattractant field. The real life difference between the two (the diffusivity of IFN is two orders of magnitude faster [E]) is immaterial to the role the fields play in the model, namely, the gradient that provides recruitment and chemotaxis signals.

**Tumor Cell Transitions**

The transition between OXPHOS and glycolytic types is determined by the amount of oxygen available to the cell and the fitness threshold (a parameter that represents the overall aerobic fitness of the patient).

$$P(\sigma(OXPHOS) \rightarrow \sigma(Glycolytic)) = TP \text{ } if \text{ } O(\sigma) < F_t \text{ (6a)}$$

$$P(\sigma(Glycolytic) \rightarrow \sigma(OXPHOS)) = TP \text{ } if \text{ } O(\sigma) > F_t \text{ (6b)}$$

Where $\sigma(OXPOS)$ represents the cell type OXPHOS, $\sigma(Glycolytic)$ represents the cell type Glycolytic, $O(\sigma)$ represents the concentration of oxygen at the center of mass of the cell. The fitness parameter $F_t$ represents the oxygen concentration at which tumor cells switch metabolic profile. Equation 6a describes the transition probability $TP$ from OXPHOS to Glycolytic when the oxygen concentration drops below the fitness threshold $F_t$. Equation 6b describes the transition probability $TP$ from Glycolytic to OXPHOS when the oxygen concentration increases above the fitness threshold $F_t$. Both OXPHOS and Glycolytic cells can transition to the Necrotic cell type if the oxygen available to the cell drops below the necrotic threshold.

$$P(\sigma(OXPOS \text{ } or \text{ } Glycolytic) \rightarrow \sigma(Necrotic)) = TP \quad if \text{ } O(\sigma) < N_t \text{ (7)}$$

Where $\sigma(Necrotic)$ represents the cell type Necrotic and $N_t$ is the oxygen concentration below which tumor cells transition to Necrotic. Tumor cells can also transition to the Apoptotic cell type if they come into contact with an active tumor suppressor immune cell.

Parameter Estimation. The transition probability was determined assuming that tumor cells do not immediately shift their metabolic profile. Instead, they persist in their current state for 2.5 hrs after the metabolic resources in their environment dropped below a certain threshold. The ranges of the fitness threshold were determined by the sensibility analysis (see *Sensitivity Analysis*). The fitness parameter is related to the experimental fitness score by the following linear relation: $1.85 * F_t + 0.08$. The necrotic threshold was determined by assuming that cells become necrotic when the oxygen tension in tissue drops below 10% of the steady state concentration of oxygen in the tissue [F].

**Cell Growth and Mitosis**

Although we do not model the cell cycle explicitly, we represent it implicitly by having tumor cells grow and divide once they have doubled their volume. We assumed that tumor cells divide at a rate that is independent of their glycolysis/OXPHOS ratio (see, e,.g., the similar KI-67 levels in both "sedentary" and "trained" tumors from [27], cited in the main article). The cell volume growth rate is a function of the local availability of oxygen, which in this case represents metabolic resources:

$$\frac{\partial V(\sigma)}{\partial t} = G\frac{O(\sigma)^2}{O(\sigma)^2 + K_O^2} \text{ (8)}$$

where $V$ is the target volume of the cell $\sigma$, $G$ is the maximum growth rate of individual tumor cells, $O$ is the concentration of oxygen at the center of mass of cell $\sigma$ and $K_O$ is the concentration of oxygen at which the growth rate decreases to half maximum. We assumed this Michaelis-Menten form for the growth rate to model a smooth transition between actively dividing and quiescent tumor cells. Although the target volume of the cell increases at a rate that is dependent only on the availability of oxygen, the actual volume of the cell depends on other factors, such as availability of space. The target surface of the cell was also adjusted such that $S(\sigma) = 4\sqrt{V(\sigma)}$, where $S$ is the new target surface of the growing tumor cell $\sigma$. The plane of division was assumed to be random.

Parameter estimation. We assumed that the average time for tumor cell division is 24 hours. Since cells must double their initial target volume to divide, we estimated the maximum growth rate of the cell to be 2.96 x 10-3 $\mu m^{2}/s$. The parameter $K_O$ is the oxygen concentration at which the growth rate is half maximum ( $\frac{G}{2}$ ) and we assume this threshold is reached when the oxygen concentration inside the tumor drops to half the steady state oxygen tension in tissue 2.15 x $10^{-4}$ mol/L. The growth rate is the same for OXPHOS and glycolytic cells.

**Chemotaxis**

Immune cells respond to chemokine signals in the TME and can actively migrate to areas of higher concentrations of such signals. In particular, IFN$\gamma$ enhances the cytolytic ability and the kinematics of Cytotoxic CD8 $^{+}$ T lymphocytes (CTL) both by paracrine and autocrine mechanisms of signaling. CTLs' search patterns in peripheral tissues are mainly dictated by informed motion in which haptotaxis and haptokinesis cues restrict their movements, and chemoattractants guide them through a signaling gradient. To model CTL migration inside the tumor, we assumed that tumor suppressors migrate against the gradient of the oxygen field, which is an indicator of highly proliferative tumor cells. We assume that both immune tumor promoters are attracted by the chemoattractant field, which drives immune cell migration to highly glycolytic areas of the TME. We also assume that immune tumor suppressors are attracted by the IFN$\gamma$ field, which drives them to areas of high density of active immune suppressors. In CPM, chemotaxis is represented as an additional energy term in the effective energy function $H$. The change in energy due to chemotaxis is calculated by considering the chemotactic force that favors voxel attempts in which cells move up or down the gradient of the field. The chemotactic force exerted over each voxel of the cells is given by:

$$F_{chem}(x) = \frac{\lambda_{f_i}}{1+f_i(\sigma)} \nabla f_i(x) \quad (9)$$

where $\lambda_{f_i}$ is the chemotactic sensitivity parameter of the cell at position $x$ to the chemical field $f_i$, $f_i(\sigma)$ is the concentration of the field at the center of mass of cell $\sigma$, and $\nabla f_i(x)$ is the gradient of the field at position $x$ with respect to the source voxel. Parameters $\lambda_{O-Supp}$ and $\lambda_{IFN-Supp}$ are the sensitivity of tumor suppressors to the oxygen and the IFN$\gamma$ fields, respectively. Parameter $\lambda_{L-Prom}$ is the sensitivity of tumor promoters to the chemoattractant field.

Parameter estimation. The chemotactic sensitivity $\lambda_{f_i}$ parameter is a CPM-specific parameter, and as such does not correspond to a measurable experimental quantity. However, since $\lambda_{f_i}$ modulates the chemotactic strength and the speed of cell migration, we chose values of -500 for and 500 for $\lambda_{IFN-Supp}$ and $\lambda_{L-Prom}$ such that immune cells migrate faster than tumor cells and penetrate the highly packed tumor while still preserving their shape and form.

**Immune Cell Recruitment**

Our model currently covers the TME only and thus ignores the complicated signaling networks behind immune cells migration, search strategies and mechanisms from the lymphoid organs to target organs (future model expansion will include these factors). We model the recruitment of Immune cells into the

TME by assigning probabilities to different immune cells appearing in the TME. In the case of a tumor suppressor immune cell, this he probability is constant:

$$\Pr(adding\ tumor\ suppressor) = R_{Supp} \quad (10)$$

Where $R_{Supp}$ is the probability per simulation unit time. To determine the seeding location of the tumor suppressor, the simulation space is randomly sampled 6 times, and a tumor suppressor immune cell is seeded at the unoccupied location with the highest amount of the IFN field. The immune cell is not seeded if none of the 6 locations is unoccupied.The probability of adding a tumor promoter immune cell depends on the fraction of glycolytic cells to all proliferative tumor cells, which we use as a proxy of the glycolytic state of the tumor.

$$\Pr(adding\ tumor\ promoter) = R_{Prom} \frac{N_{Gly}}{N_{OXP}+N_{Gly}} \quad (11)$$

Where $R_{Prom}$ is the maximum probability per simulation unit time, $N_{Gly}$ is the number of glycolytic tumor cells and $N_{OXP}$ is the number of OXPHOS tumor cells. To determine the seeding location of the tumor promoter, the simulation space is randomly sampled 6 times, and a tumor promoter immune cell is seeded at the unoccupied location with the highest amount of the chemoattractant field. The immune cell is not seeded if none of the 6 locations is unoccupied. Once the cells are in the TME, they react to their respective recruiting signals: Tumor suppressors move along the oxygen and the IFN $\gamma$ gradient (which is then reinforced once more Tumor suppressors kill tumor cells) and immune suppressors move along the chemoattractant gradient (which is modulated by changes in Glycolysis/OXPHOS balance).

Parameter estimation. The constant recruitment rate of tumor suppressors was determined such that at the end of the simulation, the density of tumor cells was 50 cells/mm $^2$. This density is 10 times smaller than the average density observed *in vivo* (from [16], cited in the main article). This scaling factor was adopted because of limitations to the spatial dimensions of the immune cells imposed by their interactions with the tumor cells (e.g. the compactness of the core of the tumor necessitated immune cells with a target volume 1.5 times bigger than the tumor cells and comparatively large to the size of immune cells *in vivo*). The recruitment rate of tumor promoters was calibrated such that by the end of the simulation, the average density of tumor suppressors was half of the density of tumor promoters, as observed *in vivo* (from [16], cited in the main article) but ranged from 1:8 to 1:1 [G].

**Immune Cell Cytotoxicity**

CTLs are found in many solid tumors and provide an attractive target for immunotherapeutic manipulation. We modeled CTL-mediated cytotoxicity by having tumor suppressors kill all the tumor cells they come in contact with. When a tumor suppressor cell comes into contact with a tumor cell, the tumor cell transitions to an apoptotic tumor cell. To model cell death, the target volume of the apoptotic cell decrease at a constant rate:

$$\frac{\partial V(\sigma)}{\partial t} = -A \quad (12)$$

where $V$ is the target volume of the cell $\sigma$, $A$ is the death rate of the apoptotic tumor cell. Since we assume that our cells occupy the space of a square, the target volume of the cell was also adjusted such that $S(\sigma) = 4\sqrt{V(\sigma)}$, where $S$ is the target surface of the apoptotic tumor cell $\sigma$. We assumed that tumor suppressors can kill more than one target tumor cell at a time.

Parameter Estimation. We assumed that a tumor cell dies after 12 hours of coming into contact with a CTL cell. This number is relatively low death rate compared to the experimental observation that apoptotic cells die within an hour of coming into contact with cytotoxic immune cells. We justify this choice of parameter value by noting that our tumor suppressor cells kill multiple cells at a time (from 1 to 6), such that the effective per CTL killing rate varies from 12 to 2 hours [H].

**Immune Cell Inhibition**

Foxp3+ T regulatory (Treg) cells are an important population of leukocytes that control immunity, mainly by dampening effector T cell responses. We assumed that CTL inhibition occurs either by contact with Trges or via inhibitory molecules secreted by Tregs (e.g., transforming growth factor (TGF)-β, IL-10, and IL-35 that bind to immune cells and result in immunosuppressive effects). We modeled the inhibitory effect of Tregs by having tumor suppressor become inhibited if they were within a given distance of a tumor promoter immune cell. We measure the distance between the center of mass of every tumor promoter cell and every tumor suppressor. If the tumor suppressor is within the inhibition radius of a tumor promoter, then it changes type to inhibited. Inhibited tumor suppressors can become active if they move away from the inhibition radius of the tumor promoter.

$$I_d \geq \sqrt{(X(\sigma_i(supp) - X(\sigma_j(prom))^2 + (Y(\sigma_i(supp) - Y(\sigma_j(prom))^2} \quad (13)$$

where $X(\sigma_i(supp))$ is the position of the center of mass of tumor suppressor cell $\sigma_i$ in the $x$ dimension, $X(\sigma_j(prom))$ is the position of the center of mass of tumor promoter cell $\sigma_j$ in the $x$ dimension, $Y(\sigma_i(supp))$ is the position of the center of mass of tumor suppressor cell $\sigma_i$ in the $y$ dimension, $Y(\sigma_j(prom))$ is the position of the center of mass of tumor promoter cell $\sigma_j$ in the $y$ dimension and $I_r$ is the inhibition radius.

Parameter Estimation. We assumed that the effect of inhibitory molecules secreted by regulatory immune cells into the TME span over six cell diameters (96 um). This assumption is justified because inhibition by T-regs occurs not only by contact, but also by secretion of inhibitory chemokines.

**Immunotherapy**

In this paper we focused on ICI (Immune Checkpoint Inhibitors) which mirror the effect of immune suppressors. For this reason, and as recently shown *in vivo* (see main paper), we assumed a tradeoff between immune suppressors density in the TME (which is a function of aerobic fitness, according to our hypothesis) and ICI dosage, and, as a result, a tradeoff between aerobic fitness and potential adverse side effects of ICI (measured in our model with cytotoxicity levels), when the dosage is personalized to the subject's aerobic fitness (the more aerobically fit the subject, the lower dosage of ICI they can be given, with similar tumor reduction outcomes relative to a sedentary subject, but with lower cytotoxicity. We model the mechanism of action of ICI dosage by reducing the inhibition radius of tumor promoters. ICI dosages are represented as different magnitudes of the reduction in the inhibition radius. ICI-associated cytotoxicity is measured as the additional exposure (area under the curve) to IFN$\gamma$ produced by tumor suppressors with respect to the baseline (no treatment).

### Parameters

| Conversion Factors | | |
|---|---|---|
| MCS | 360.0 s | |
| Lattice length | 4.0 um | |
| Concentration conversion factor | 1.0 X 10 16 mol | |
| Parameter | Symbol | Value |
| Initial target volume tumor cell (Eq 1) | $V_T$ | 256 um^2 |
| Lambda volume (Eq 1) | $\lambda_{vol}$ | 16 |
| Initial target surface tumor cell | $S_T$ | 64 um |
| Lambda surface | $\lambda_{sur}$ | 16 |
| Target volume immune cell (Eq1) | $V_T$ | 384 um^2 |
| Lambda volume (Eq 1) | $\lambda_{vol}$ | 24 |
| Target surface immune cell | $S_T$ | 78.4 um |
| Lambda surface | $\lambda_{sur}$ | 19.6 |
| Membrane fluctuation (Eq 2) | $T$ | 50.0 |
| Oxygen diffusion coefficient (Eq 3) | $D_O$ | 1460.0 um^2/s |
| Oxygen uptake by OXPHOS tumor cells (Eq 3) | $d_{OXPHOS}$ | 6.00 x 10-17 mol/(cell*s) |
| Global oxygen decay rate (Eq 3) | $d_{global}$ | 0.01 1/s |
| Oxygen production by Medium (Eq. 3) | $P_{Medium}$ | 8..00 x 10-16 mol/s |
| Chemoattractant diffusion coefficient (Eq 4) | $D_L$ | 0.1 um^2/s |
| Chemoattractant decay rate (Eq 4) | $d_L$ | 1.0 x 10-7 1/s |
| Chemoattractant production by glycolytic c cells (Eq 4) | $P_{L-Gly}$ | 7.52 x 10-17 mol/(cell*s) |
| IFN-γ diffusion coefficient (Eq 5) | $D_{IFN}$ | 0.1 um^2/s |
| IFN-γ decay rate (Eq 5) | $d_{IFN}$ | 1.0 x 10-7 1/s |

| | | |
|---|---|---|
| IFN-γ production by tumor suppressor cells (Eq 5) | $P_{IFN-Immune}$ | 7.52 x 10-17 mol/(cell*s) |
| Fitness Threshold (eq 6a,b) | $F_t$ | [2.34x10^-4 to 9.38x10^-5] mol/L] |
| Necrotic Threshold (eq 7) | $N_t$ | 6.25 x 10-5 mol/L |
| Cell Transition Probability (eqs 6-7) | $TP$ | 0.000111 1/s |
| Maximum tumor growth rate (Eq 8) | $G$ | 2.96 x 10-3 um^2/s |
| Oxygen concentration at which tumor growth rate is half maximum (Eq 9) | $K_O$ | 2.15 x 10 -4 mol/L |
| Chemotaxis sensitivity of tumor suppressors to oxygen field (Eq 10) | $\lambda_{O-Supp}$ | -500 |
| Chemotaxis sensitivity of tumor suppressors to IFN field (Eq 10) | $\lambda_{IFN-Supp}$ | 500 |
| Chemotaxis sensitivity of tumor promoters to chemoattractant field (Eq 10) | $\lambda_{L-Prom}$ | 500 |
| Recruitment rate of tumor suppressor immune cells (Eq 11) | $R_{Supp}$ | 0.0111 cell/s |
| Recruitment rate of tumor suppressor immune cells (Eq 11) | $R_{Prom}$ | 0.0167 cell/s |
| Apoptosis rate (Eq 12) | $A$ | 3.70 x 10-4 um^2/s |
| Tumor promoter inhibition radius (Eq 13) | $I_d$ | 96 um |

### Sensitivity Analysis - Fitness Parameter

| Fitness Parameter Pair | p-value |
|---|---|
| 0.05 - 0.06 | 0.7376007368052694 |
| 0.06 - 0.07 | 0.17069549494379443 |
| 0.07 - 0.08 | 0.13780438758905694 |
| 0.08 - 0.09 | 0.09456554045876897 |
| 0.09-0.10 | 0.23643686583200896 |
| 0.10-0.11 | 0.11812682730957082 |
| 0.11-0.12 | 0.34610341421792135 |
| 0.12-0.13 | 0.21273526017530378 |
| 0.13-0.14 | 0.10195752002748157 |
| 0.-14-0.15 | 0.03583599939175715 |
| 0.15-0.16 | 0.41735667350462025 |

### Standard Error - Number of Replicates

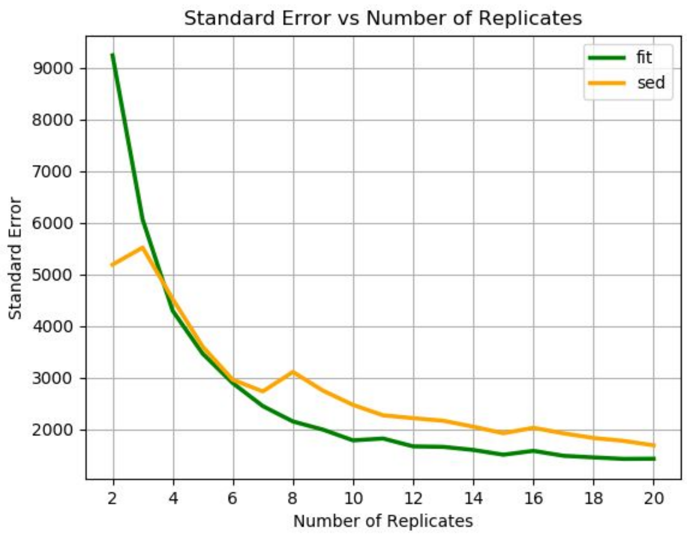


Figure S1: Standard Error vs Number of Replicates. The standard error is minimized after 20 simulation replicates.

### Additional Plots

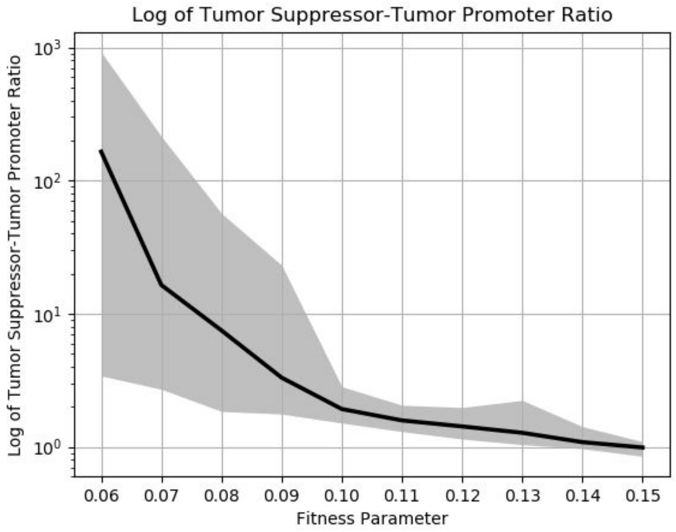


Figure S2: Tumor suppressor- Tumor Promoter ratio vs Fitness Parameter. The ratio of tumor suppressors to tumor promoters decreases exponentially with increase in the fitness parameter.

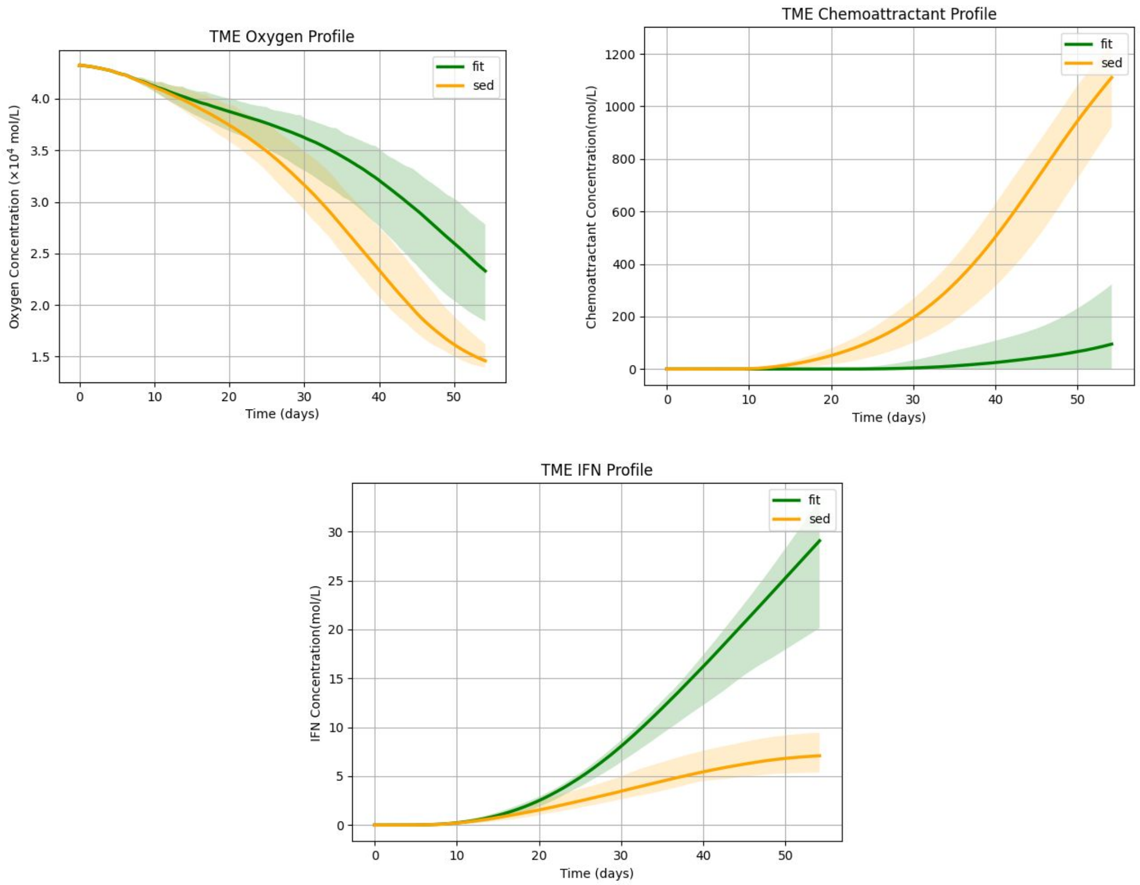

Figure S3: Chemical fields concentrations vs time for fit and sedentary groups. The oxygen profile in the TME is lower in the sedentary group because of the higher number of tumor cells. The chemoattractant profile is higher in the sedentary group because of the higher proportion of glycolytic cells. The IFN profile is higher in the fit group because of the higher number of active tumor suppressors.

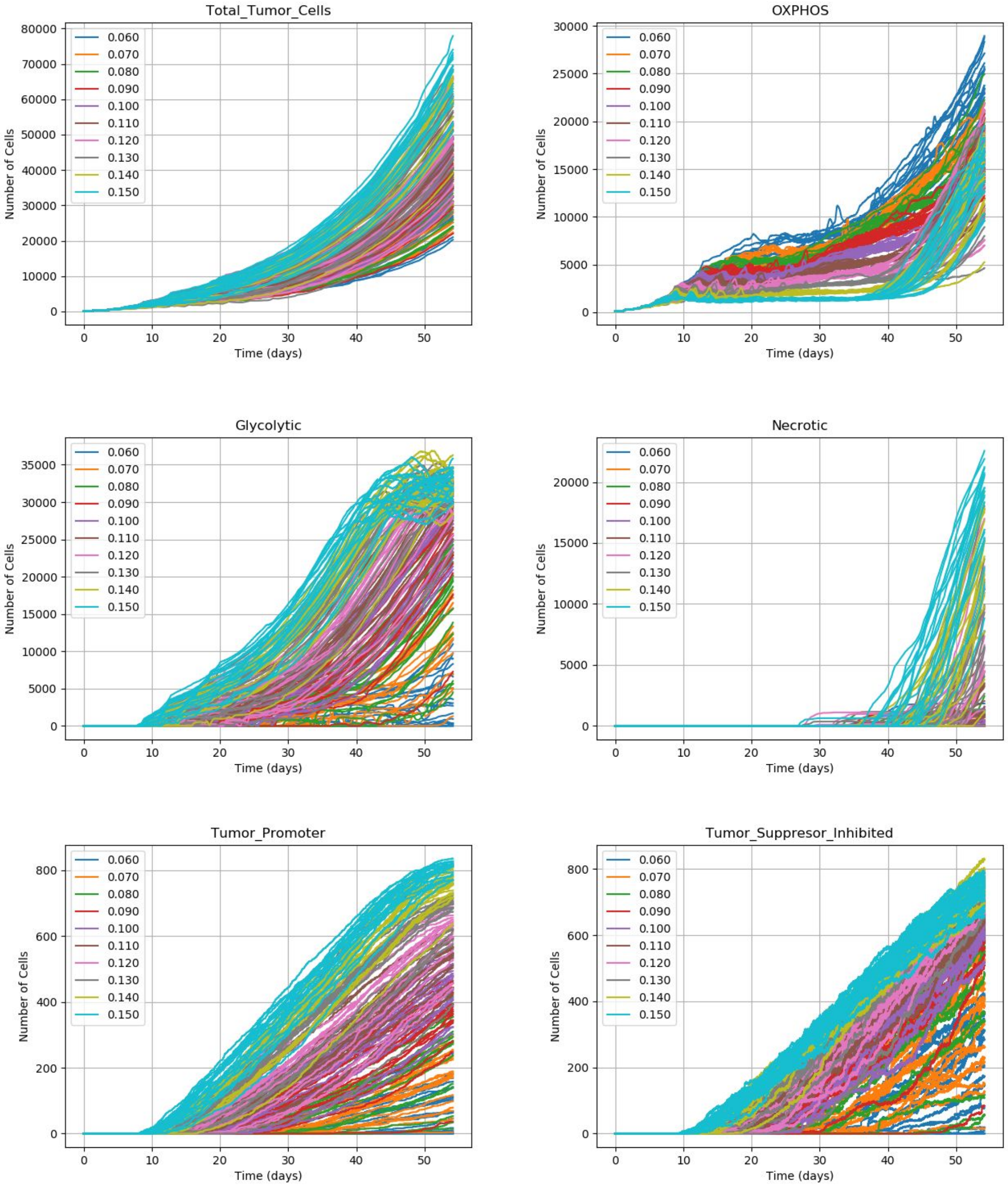

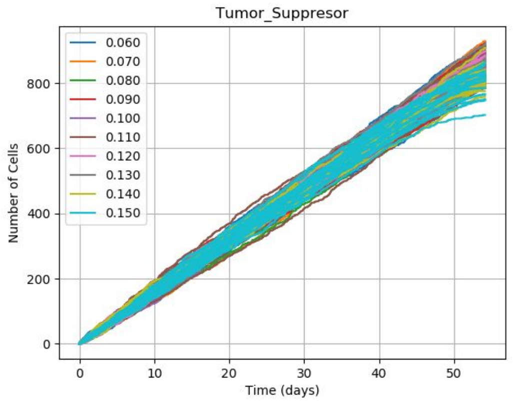


Figure S4: Number of cells for individual simulation replicates by fitness group.

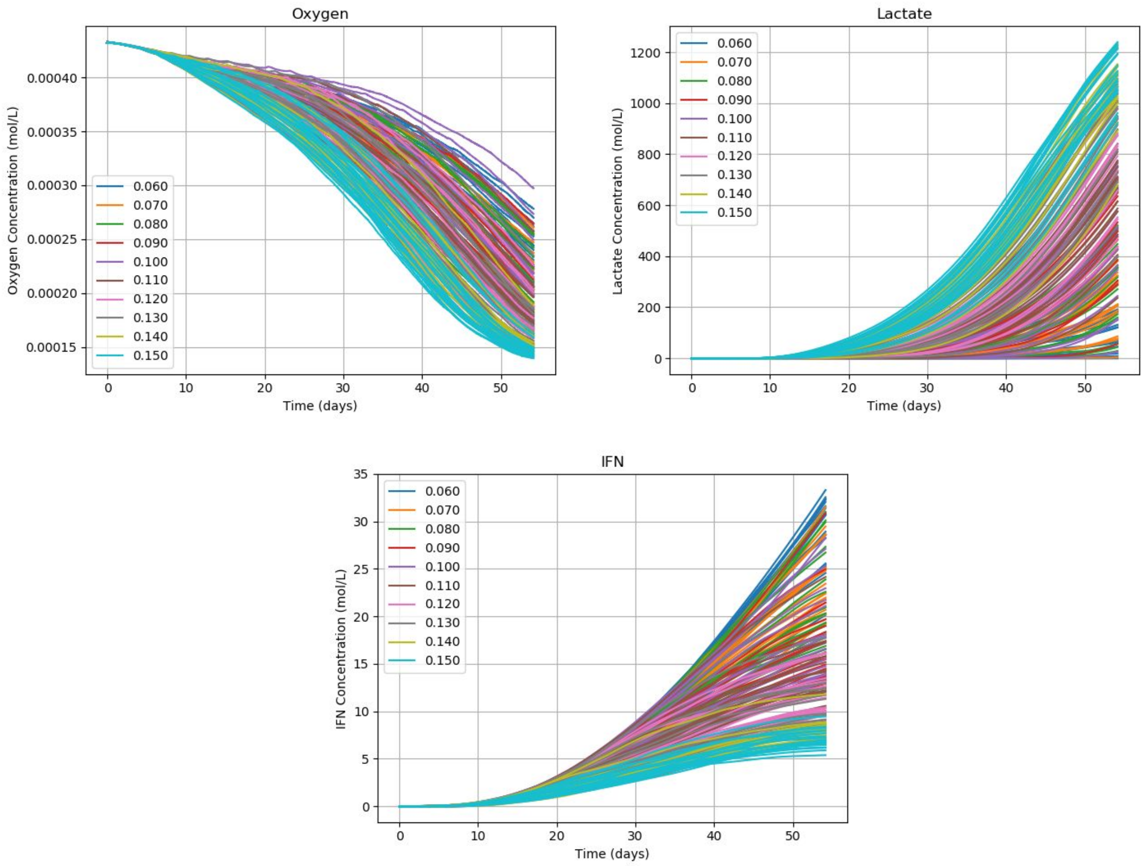


Figure S5: Chemical fields concentration for individual simulation replicates by fitness group.
